# High-speed quantitative optical imaging of absolute metabolism in the rat cortex

**DOI:** 10.1101/786244

**Authors:** R. H. Wilson, C. Crouzet, M. Torabzadeh, A. Bazrafkan, N. Maki, B. J. Tromberg, Y. Akbari, B. Choi

## Abstract

Quantitative measures of blood flow and metabolism are essential for improved assessment of brain health and response to ischemic injury. In this report, we demonstrate a multimodal technique for measuring the cerebral metabolic rate of oxygen (CMRO_2_) in the rodent brain on an absolute scale (μM O_2_ / min). We use laser speckle imaging (LSI) at 809 nm and spatial frequency domain imaging (SFDI) at 655 nm, 730 nm, and 850 nm to obtain spatiotemporal maps of cerebral blood flow (CBF), tissue absorption (μ_a_), and tissue scattering (μ_s_’). Knowledge of these three values enables calculation of a characteristic blood flow speed, which in turn is input to a mathematical model with a “zero-flow” boundary condition to calculate absolute CMRO_2_. We apply this method to a rat model of cardiac arrest (CA) and cardiopulmonary resuscitation. With this model, the zero-flow condition occurs during entry into CA. The CMRO_2_ values calculated with our method are in good agreement with those measured with magnetic resonance (MR) and positron emission tomography (PET) by other groups. Our technique provides a quantitative metric of cerebral metabolism that can potentially be used for comparison between animals and longitudinal monitoring of a single animal over multiple days, to assess differences in baseline metabolism and track recovery of metabolism in survival studies following ischemia and reperfusion.

## Introduction

Assessing brain metabolism on a quantitative scale is critical for improved diagnosis, monitoring, and treatment of ischemic damage caused by conditions such as cardiac arrest (CA) and focal stroke. In particular, it is critical to measure the *absolute* cerebral metabolic rate of oxygen (CMRO_2_), Such knowledge would obviate the need for baseline measurements and facilitate longitudinal measurements to track longer-term cerebral recovery following ischemia and reperfusion. Also, it is essential to measure absolute CMRO_2_ to enable quantitative comparisons between the values of different subjects at baseline and at subsequent time points in preclinical or clinical studies. These needs are currently unmet, as no existing technology directly and noninvasively measures absolute CMRO_2_ with high spatial and temporal resolution using endogenous contrast.

Conventional clinical monitoring techniques (e.g., arterial blood pressure, jugular bulb oximetry, pulse oximetry, laser Doppler flowmetry) are typically unable to separate alterations in cerebral metabolism from changes in blood flow. Measurement of CMRO_2_ requires medical imaging techniques such as Positron Emission Tomography (PET) or functional Magnetic Resonance Imaging (fMRI). PET [1-5] can measure absolute CMRO_2_, but it is expensive, non-portable, and uses exogenous contrast agents containing radioactive tracers. fMRI measures CMRO_2_ changes via the blood oxygen level dependent (BOLD) signal, but it struggles to provide absolute CMRO_2_ without extensive calibration because it uses the BOLD signal as a surrogate for cerebral blood flow and hemoglobin content as opposed to directly measuring these quantities. Additionally, both PET and BOLD fMRI typically have limited temporal resolution and cannot be performed repeatedly on a patient over a short period.

Diffuse optical imaging (DOI) techniques are an inroad to addressing this problem, as they are noncontact and use non-harmful visible and near-infrared light to probe tissue absorption and scattering for endogenous contrast. Recently, we demonstrated that the combination of Spatial Frequency Domain Imaging (SFDI) and Laser Speckle Imaging (LSI)) can quantify tissue metabolic changes with high spatial and temporal resolution [6]. We also recently applied high-speed LSI [7] and SFDI [8] to measure perfusion, oxygenation, and tissue scattering in the brain in a cardiac arrest (CA) model of global cerebral ischemia. The capability of our rapid multimodal SFDI+LSI system to image blood flow and hemoglobin concentration simultaneously enables high-speed measurement of CMRO_2_ [9].

Previous optical methods to calculate CMRO_2_ have typically used point source techniques [10] and intrinsic signals that do not correct for changes in tissue scattering [9, 11]. However, by combining LSI and multispectral SFDI, we can account for the effects of time-varying tissue scattering at multiple wavelengths [12] and the contribution of venous regions versus parenchyma when calculating CMRO_2_. Furthermore, most optical techniques measure only the relative CMRO_2_, as they are unable to solve for the necessary quantities to calculate the absolute CMRO_2_. Therefore, these methods typically require measurement of a baseline before each imaging session, so that changes in CMRO_2_ during the experiment can be assessed relative to that baseline. This approach makes it very difficult to perform quantitative longitudinal studies of metabolic changes over periods of days or weeks where multiple separate imaging sessions are required. By contrast, the combination of our multimodal instrumentation and CA model enables *continuous measurement of absolute CMRO*_*2*_ during the dynamic cerebral ischemia and reperfusion phenomena caused by CA and resuscitation.

Here, we report a technology that uses multimodal optical imaging (LSI + SFDI), along with a mathematical model of CMRO_2_ that incorporates a “zero-flow” boundary condition during the onset of ischemia in our CA model, to obtain the parameters necessary for absolute CMRO_2_ measurement. By merging quantitative measurement of absolute CMRO_2_ with high-speed resolution (up to ∼14 Hz) of rapid changes in CMRO_2_ during highly dynamic events (e.g., CA and reperfusion), our approach represents a significant potential step forward in quantitative metabolic imaging of the brain.

## Methods

### Animal Preparation

The Institutional Animal Care and Use Committee (IACUC) at the University of California, Irvine (protocol number 2013-3098-01) approved all of the procedures described in this report. Ten male Wistar rats (weight ∼300-400 g) were imaged for this study, and the animal preparation procedures are described in our previous publications [7, 8, 13, 14]. Before the experiment, all rats were endotracheally intubated under isoflurane anesthesia. Each rat had epidural screw electrodes implanted for electrocorticography (ECoG). Each rat was given a hemicraniectomy (4 mm right-to-left × 6 mm anterior-to-posterior), which enabled imaging of a portion of the right sensory and visual cortex. Cannulation of the femoral artery was performed to allow the delivery of drugs, the sampling of blood, and the monitoring of blood pressure.

### Cardiac Arrest (CA) and Cardiopulmonary Resuscitation (CPR)

Fig. 1 shows the multimodal setup employed in the experiments. At the onset of each experiment, the level of isoflurane was decreased from 2% to 0.5-1%. Concurrently, the mixture of inhaled gases was altered from 50% O_2_ + 50% N_2_ to 100% O_2_. Two minutes later, to reduce confounding effects of isoflurane on cerebral perfusion and metabolism, the anesthesia was turned off entirely after an additional two minutes, at which time the rat was exposed to room air (21% O_2_). During this same time period, 1 mL of 2 mg/kg Vecuronium (a neuromuscular blocker) and 1 mL of heparinized saline were administered intravenously, which enabled respiration to be controlled entirely by the ventilator. This stage of the experiment lasted for 3 min, after which the ventilator was turned off to induce asphyxia and cause progressive hypoxic hypercarbic hypotension. CA was defined as the period over which the pulse pressure decreased below 10 mmHg and systolic pressure below 30 mmHg. The conditions of these experiments induced pulseless electrical activity, which is common in CA patients in a hospital setting.

**Fig. 1.**
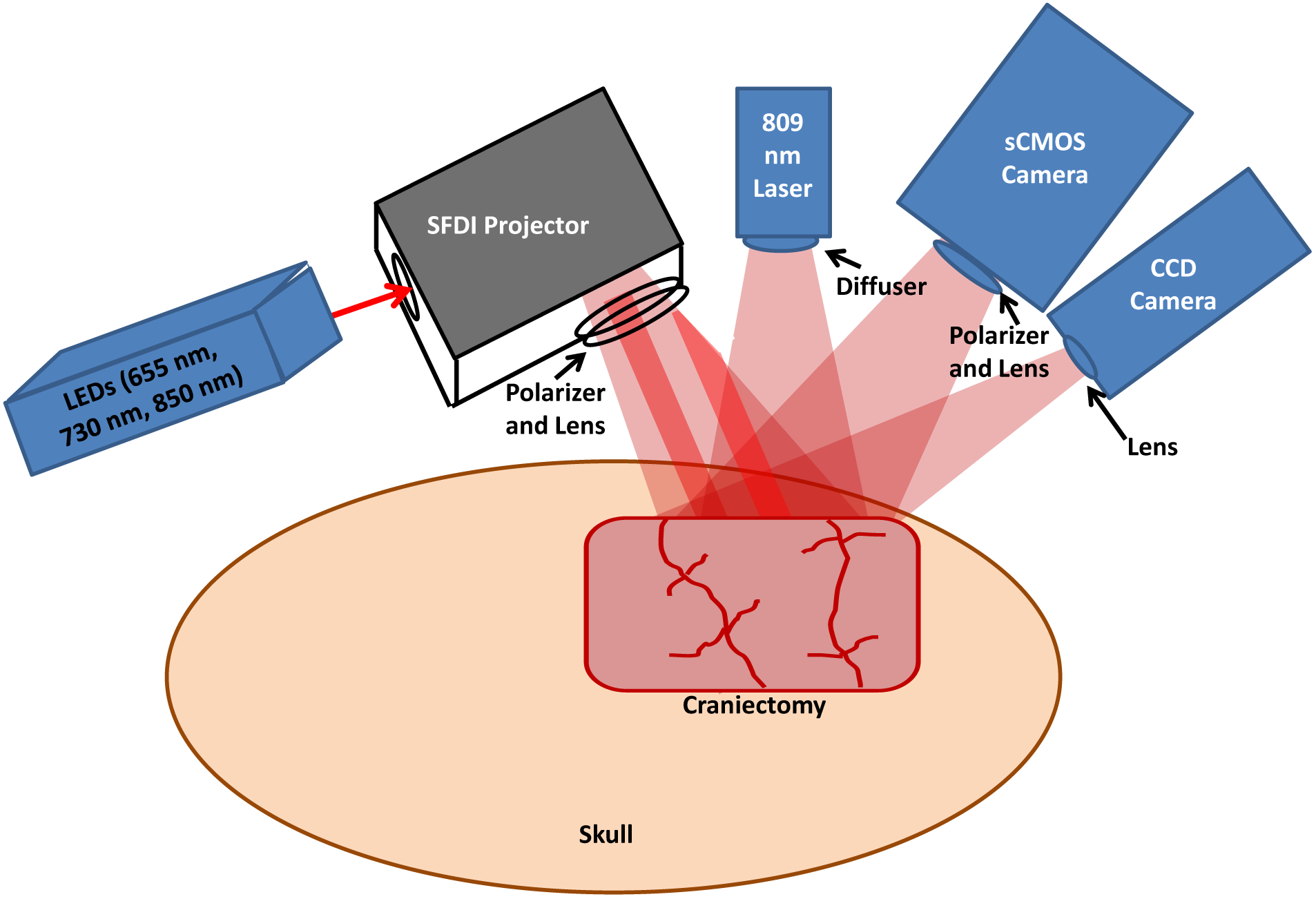
Multimodal platform (not to scale) for Laser Speckle Imaging (LSI) and multispectral Spatial Frequency Domain Imaging (SFDI) of the rat brain. A craniectomy (∼6 mm × 4 mm area) is performed to provide direct access to the brain for optical imaging. For SFDI, light-emitting diodes (LEDs) of 655 nm, 730 nm, and 850 nm are sequentially sent into a spatial light modulator that acts as a projector to send spatially-modulated patterns of light onto the brain [8]. A scientific CMOS camera detects the backscattered light. For LSI, an 809 nm laser illuminates the brain with coherent light, and the remitted speckle pattern is captured at 60 fps with a CCD camera.

Forty-five seconds before onset of CPR, the ventilator was turned back on (respiratory rate = 85 breaths/min, PIP = 17.5-18.5 cmH_2_O, PEEP = 3 cmH_2_O at 2.5 LPM), and 100% oxygen was given. Immediately before the onset of CPR, 0.01 mg/kg epinephrine, 1 mmol/kg sodium bicarbonate, and 2 mL of heparinized saline were administered intravenously. Then, CPR was performed via external cardiac massage. CPR was terminated upon return of spontaneous circulation (ROSC), as identified from arterial blood pressure measurements. Subsequently, the animal was monitored continuously with arterial blood pressure, LSI, SFDI, and ECoG for an additional ∼2 hr, after which the animal was euthanized with pentobarbital. Recovery of ECoG signal following ROSC was quantified by (1) time to initial resumption (burst) of ECoG activity, and (2) ECoG Information Quantity (IQ) 90 min post-ROSC (as in [15]).

### Laser Speckle Imaging (LSI)

For LSI, an 809 nm laser with long coherence length (Ondax, Monrovia, CA) served as the light source. To make the illumination close to uniform, we placed a ground-glass diffuser (ThorLabs, Inc., Newton, NJ) between the laser and the tissue. A CCD camera (Point Grey Research Inc., Richmond, BC, Canada) detected the backscattered light with a 10 ms exposure time, enabling image acquisition at a frame rate of 60 Hz. Using a 5×5 sliding spatial window filter, the equation K = σ/<I> was employed to calculate the local speckle contrast K at each pixel, where <I> was the mean intensity over the window and σ the standard deviation over the window [16]. Then, the speckle flow index (SFI) was determined from the speckle contrast K and the exposure time T via a simplified speckle imaging equation SFI = 1/(2TK^2^) [16]. Time-resolved SFI curves were generated by taking the mean of the SFI over a selected region of interest (ROI) at each time point.

### Spatial Frequency Domain Imaging (SFDI)

For SFDI, light-emitting diodes (LEDs) of three different wavelengths (655 nm, 730 nm, 850 nm) were used as light sources. The light from the LEDs was directed to a spatial light modulator that projected square-wave patterns onto the brain [8]. Backscattered light was captured using a scientific complementary metal-oxide semiconductor (sCMOS) camera (Hamamatsu Photonics). An Arduino board was used to synchronize the camera and LEDs so that images of each spatial pattern and wavelength were obtained sequentially. For each wavelength, four patterns were projected onto the tissue in sequence. The first pattern was non-modulated, and the three subsequent patterns were modulated at spatial frequency ∼0.3/mm with three distinct spatial phases to enable demodulation [17]. Thus, there were a total of (3 wavelengths × 4 frames) = 12 frames in each sequence. At the spatial frequency of ∼0.3/mm, the detected square wave pattern could be approximated as a sinusoid, allowing demodulation in the manner described by our group previously [18]. A full sequence of frames was needed to reconstruct tissue hemodynamics and CMRO_2_, resulting in an effective imaging rate of ∼14 Hz.

After demodulating the spatially-modulated data, the diffuse reflectance at each time point and wavelength was calculated from the raw data via calibration against a tissue-simulating phantom with known optical properties [17]. The diffuse reflectance was then fit with a Monte Carlo model to extract the tissue absorption coefficient μ_a_ and reduced scattering coefficient μ_s_’ [8]. Next, to improve the clarity of the μ_a_ maps, the average μ_s_’ was determined for a selected ROI and a new μ_a_ was determined using diffuse reflectance with the non-modulated pattern and this average μ_s_’. To calculate the concentrations of oxygenated and deoxygenated hemoglobin (ctHbO_2_ and ctHb, respectively) within the tissue, this new μ_a_(λ) spectrum was fit with the model spectrum μ_a_(λ) = 2.303(ctHbO_2_eHbO_2_ + ctHbεHb). The total tissue hemoglobin concentration (ctHb_tot_) was calculated by summing ctHb and ctHbO_2_. The tissue oxygen saturation was determined using the equation StO_2_ = ctHbO_2_/(ctHbO_2_ + ctHb).

### Correction of speckle flow index for tissue absorption and scattering

To correct the measured SFI for tissue absorption and static scattering, the measured speckle contrast *K* was converted to a characteristic flow speed (*v*_*c*_) by using the following equation [19]:

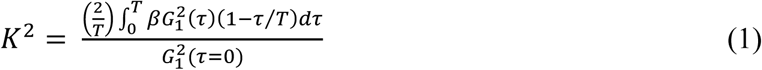

In this equation, β is a constant (typically set to 1) related to polarization and coherence properties of the LSI instrumentation. From the Siegert relationship, the intensity autocorrelation function G_2_(τ) can be related to G_1_(τ), which, in turn, is described by the correlation diffusion equation:

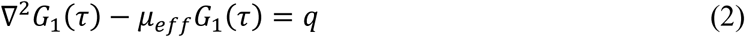

In Eq. (2), *q* is the source term and μ_eff_ = (3μ_a,dyn_μ_tr_)^1/2^, where μ_tr_ = (μ_a_ + μ_s_’) is the tissue transport coefficient and μ_a,dyn_ = (μ_a_ + μ_s_’k ^2^<Δr^2^(τ)>/3) is the dynamic tissue absorption coefficient [19]. In the equation for μ_a,dyn_, <Δr^2^(τ)> is the mean square displacement of the moving scatterers (i.e., the red blood cells) and k_o_ is the photon wavenumber. Solving Eq. (2), G_1_(τ) can be written as:

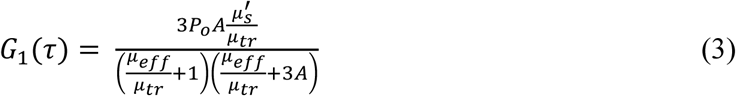

In Eq. (3), P_o_ is the incident optical power, and A is a function of the tissue refractive index [17]. All other terms in Eq. (3) are exclusively functions of the static and dynamic tissue absorption and scattering coefficients (μ_a_, μ_s_’, μ_a,dyn_). Since μ_a,dyn_ is a function of <r^2^(τ)>, and <r^2^(τ)> can be related to the characteristic flow speed *v*_*c*_ via the equation <r^2^(τ)> = *v*_*c*_τ^2^ (for directional flow) [20]. Using this framework and inputting the measured value of *K* from LSI and the measured μ_a_ and μ_s_’ from SFDI at each time point, Eq. (1) was solved for *v*_*c*_ at each time point and each pixel by iterating over a pre-defined grid of potential *v*_*c*_ values and minimizing a least-squares cost function. The resulting spatiotemporal values of *v*_*c*_ were used in place of *SFI* in the subsequent steps to achieve an optical property-corrected calculation of CMRO_2_.

### Absolute cerebral metabolic rate of oxygen (CMRO_2_) calculation

To calculate absolute CMRO_2_, we start from the equation [21]:

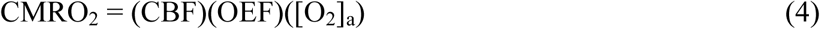

In Eq. (4), CBF is the cerebral blood flow, [O_2_]_a_ is the arterial concentration of oxygen, and OEF is the oxygen extraction fraction, equal to ([O_2_]_a_-[O_2_]_v_)/[O_2_]_a_, where [O_2_]_v_ is the venous concentration of oxygen. For a single arteriole, (OEF)([O_2_]_a_) represents the concentration of oxygen that was extracted from that arteriole and used by the brain for metabolic processes related to the synthesis of ATP. This quantity is equivalent to the concentration of deoxygenated hemoglobin that arrived in a venule following oxygen extraction by the brain. Therefore, within our measurement paradigm [6], Eq. (4) is re-written as:

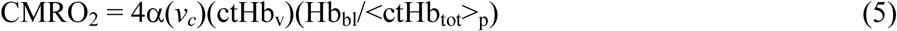

In Eq. (5), ctHb_v_ is the tissue concentration of deoxygenated hemoglobin in a region of interest atop a large vein in the ctHb maps obtained from SFDI. The factor of 4 accounts for the fact that the hemoglobin molecule has four binding sites for oxygen. Since *v*_*c*_ is still a characteristic flow parameter and not an absolute value of blood flow velocity, it is necessary to include the proportionality constant α in the equation to convert *v*_*c*_ into a quantity with units of absolute flow speed.

The factor (Hb_bl_/<ctHb_tot_>_p_) accounts for partial-volume effects caused by the diffuse nature of light propagation in the brain. Eq. (4) requires an *intra-vascular* oxygen concentration, but we measure a *bulk tissue* deoxyhemoglobin concentration. Hence, a blood-volume fraction term is required to convert between these two quantities. The numerator, Hb_bl_, is the concentration of hemoglobin in the blood sampled from the femoral artery of the animal during the arterial blood gas measurement (ABG). The denominator, <ctHb_tot_>_p_, is the mean total tissue hemoglobin concentration in the parenchyma during the period that the ABG was acquired. The factor (Hb_bl_/<ctHb_tot_>_p_) enables the required conversion of our optical ctHb measurements from the scale of a tissue hemoglobin concentration to the scale of a vascular hemoglobin concentration, removing the partial volume effect and allowing us to measure CMRO_2_ on an absolute scale.

The parameter α is a typically unknown constant; thus, the quantity reported in optical brain imaging studies is usually the relative CMRO_2_ (rCMRO_2_), since the α factor can be eliminated when calculating rCMRO_2_. However, in this report, we were able to measure *absolute* CMRO_2_ by using a “zero-flow” boundary condition [22], which is provided by the onset of cerebral ischemia in our animal model:

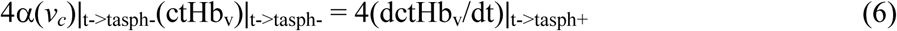

This procedure was performed for each of the 10 rats in this study, using the values of SFI and ctHb_v_ during the period immediately before asphyxia (t->t_asph_ ^−^) and the mean rate of change dctHb_v_/dt immediately after the onset of asphyxia (t->t_asph_ ^+^). The values of α and absolute CMRO_2_ then were calculated over the entire craniectomy region at each measurement time point. To our knowledge, this report represents the first high-speed diffuse optical imaging measurement of absolute CMRO_2_ in the brain with high temporal resolution during a preclinical study.

## Results

Fig. 2 shows the importance of using an ROI over a prominent vein (as opposed to an ROI in the parenchyma) for the absolute CMRO_2_ calculation. Specifically, the rate of change in cerebral ctHb during induction of the zero-flow condition (i.e., the start of asphyxia) was 58.0 ± 39.2 μM/min in the venous ROI, but only 29.7 ± 12.1 μM/min in the parenchyma. Therefore, using a parenchymal ROI or a “global” (venous + parenchyma) ROI is expected to reduce the value of α calculated with Eq. 6, leading to systematic underestimation of CMRO_2_.

**Fig. 2.**
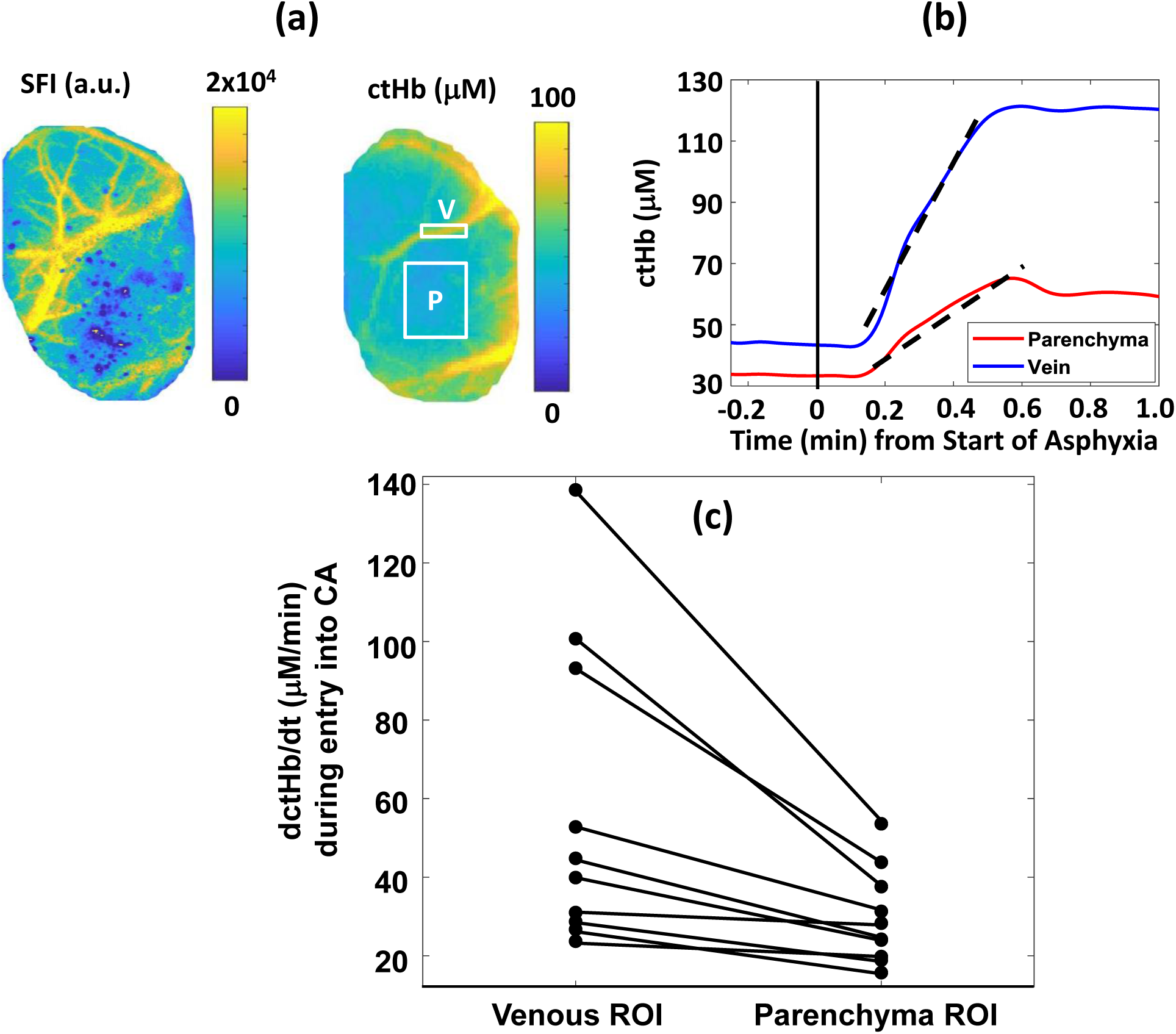
(a) Images of blood flow (speckle flow index; SFI) and deoxyhemoglobin concentration (ctHb), measured in the rat brain using LSI and SFDI, respectively. Regions of interest (ROIs) over the parenchyma (P) and large vein (V) are labeled in the ctHb image. (b) Time courses of ctHb in these ROIs during the initial min of asphyxia for the rat shown in (a), with the linear portion of the curves (used to calculate the parameter α in Eq. 6) denoted by dashed black lines. (b, c) The rate of change of tissue deoxyhemoglobin concentration (dctHb/dt) during the beginning of asphyxia is higher in the venous ROI than parenchyma ROI, demonstrating the importance of using the venous ROI for the absolute CMRO_2_ calculation.

Maps of absolute CMRO_2_ throughout a representative CA/CPR experiment are shown in Fig. 3. At baseline, the animal is under anesthesia (2% isoflurane). After 2 min of anesthesia washout, the CMRO_2_ increases by a factor of ∼2 as the animal is waking up. Following the onset of ischemia, the CMRO_2_ drops precipitously as the animal enters CA. After resuscitation, the CMRO_2_ rapidly increases until reaching a maximum at ∼8 min post-CPR (during hyperemia). Subsequently, the CMRO_2_ drops back toward baseline as ECoG bursting begins (∼12 min post-CPR).

**Fig. 3.**
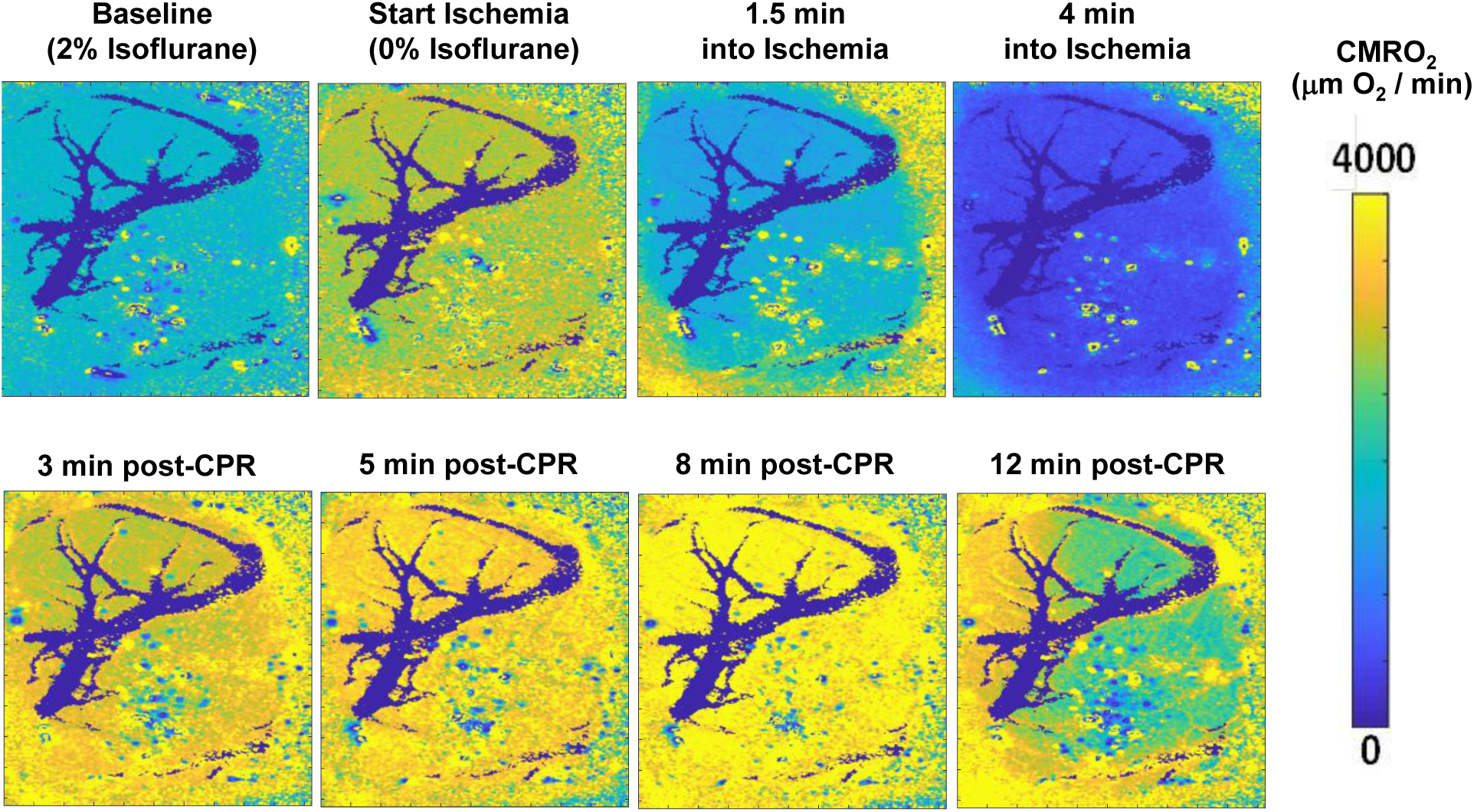
Absolute CMRO_2_ (μM O_2_/min) maps of a ∼ 6 mm × 4 mm region of the rat brain at different time points during a CA/CPR experiment. Metabolic activity increases as anesthesia is being washed out (between Baseline and Start Ischemia), followed by a sharp decrease during entry into CA (Ischemia; two rightmost top panels). Following CPR, CMRO_2_ recovers to anesthesia-free baseline level (∼3 min post-CPR), subsequently increases to levels higher than baseline (∼5-8 min post-CPR), and then declines to values approaching anesthetized baseline level once cerebral electrical activity resumes (12-min post-CPR). Large vessels (dark blue) have been removed from the CMRO_2_ images to signify that the hemoglobin in those vessels has already undergone oxygen consumption by the bulk tissue.

Fig. 4 demonstrates the effect of using the *μ*_*a*_ and *μ*_*s*_*’* from the SFDI data, along with Eq. (1), to determine a characteristic flow speed (*v*_*c*_) value that takes optical properties into account, and then using that value in place of SFI for the rCMRO_2_ calculation. During each of the experimental phases, a comparison of rCMRO_2_ trends suggests that the metabolic activity is at times overestimated (i.e., during the hyperemic phase) and underestimated (i.e., during cardiac arrest) with use of SFI instead of *v*_*c*_. This result demonstrates the effects that hyperdynamic changes in cerebral optical properties can have on calculations of CMRO_2_ dynamics using SFI alone to measure flow.

**Fig. 4.**
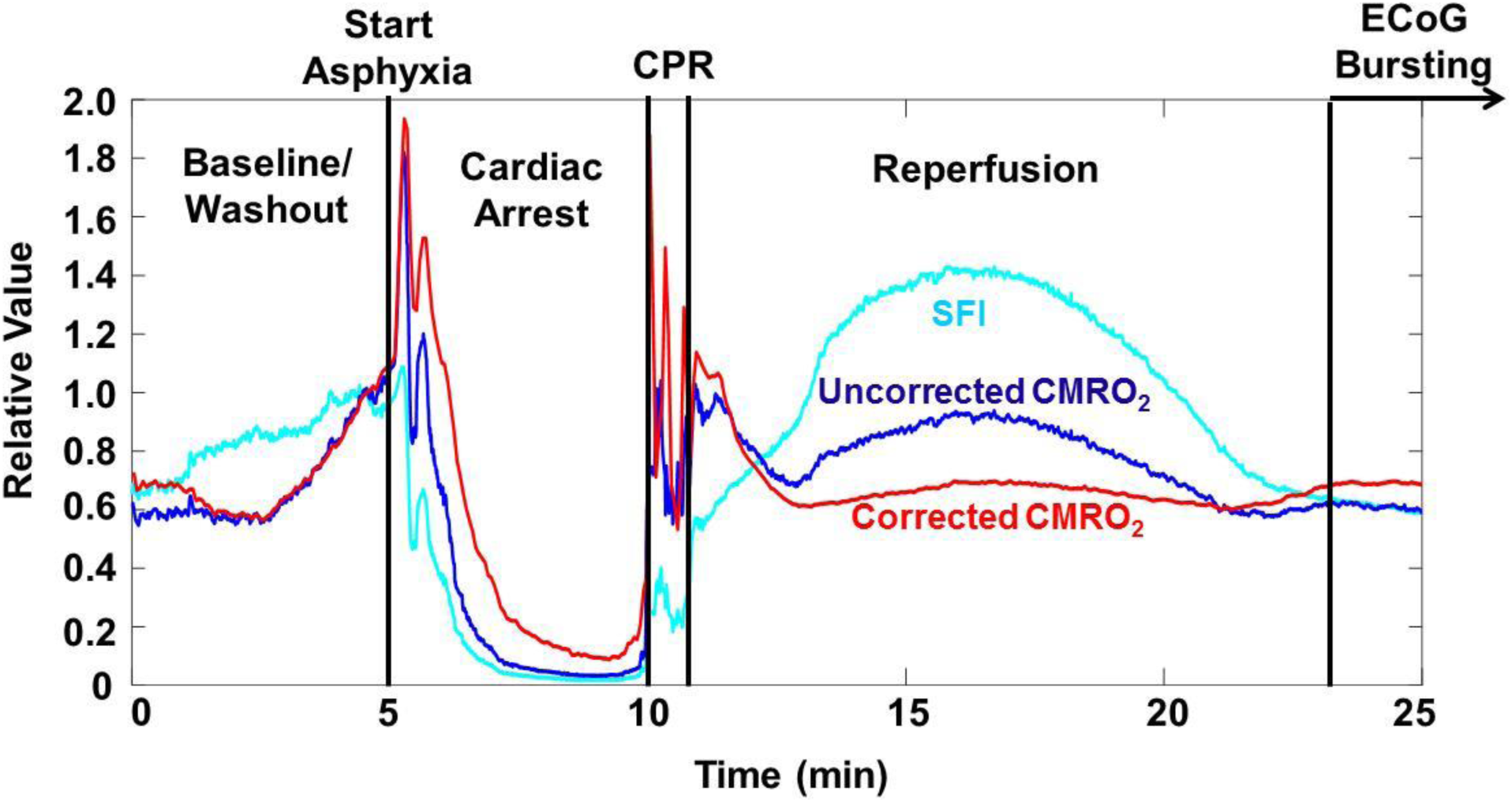
Comparison of SFI (light blue), CMRO_2_ calculated using SFI (dark blue), and CMRO_2_ calculated using *v*_*c*_ (red) after correction of SFI for tissue optical properties measured with SFDI. For ease of comparison, all three curves are normalized to their value at a point near the end of the washout period (t∼4 min). This correction reveals differences in the observed rates of change in CMRO_2_ during reperfusion and resumption of ECoG bursting, demonstrating the need to take into account optical properties even for rCMRO_2_ measurements.

Representative time courses of absolute CMRO_2_ values for regions of interest in the parenchyma of two representative rats are shown in Fig. 5(a). Interestingly, the rat with the higher baseline CMRO_2_ values required less time to resume ECoG activity (bursting) following CPR and had a higher ECoG IQ 90 min post-ROSC. For the 10 subjects included in this study, we observed that, regardless of CA duration, higher absolute CMRO_2_ values at baseline correlated significantly with cerebral electrical activity (ECoG IQ) 90 min post-CPR (Spearman; r = 0.75; p = 0.026) (Fig. 5(b)). These data collectively suggest that the absolute CMRO_2_ measurement *at baseline* may be predictive of cerebral electrical recovery *following* CA and CPR.

**Fig. 5.**
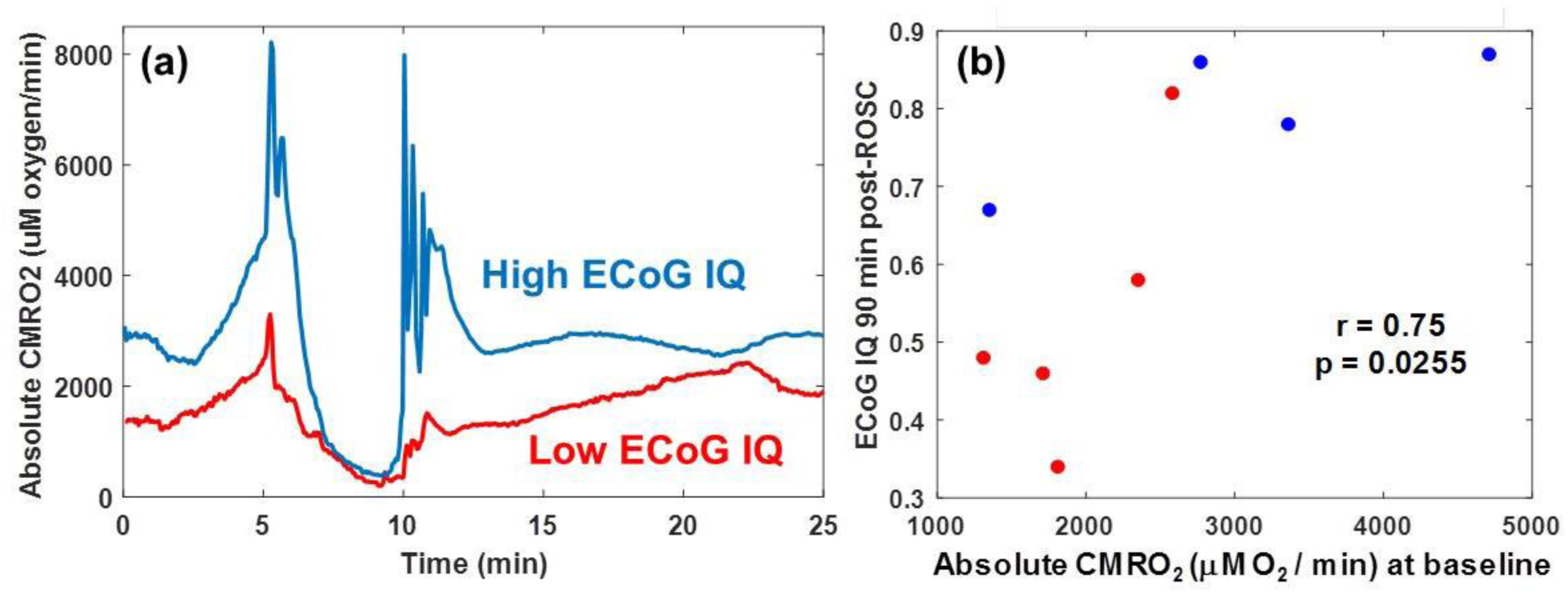
(a) Time courses of absolute CMRO_2_ for two rats. The rat with the lower initial absolute CMRO_2_ (red) required more time for resumption of ECoG bursting than the rat with the higher initial absolute CMRO_2_. In addition, 20 min after the first ECoG burst, the rat with the lower initial absolute CMRO_2_ exhibited an ECoG burst frequency ∼7x greater than the rat with the higher initial absolute CMRO_2_. (b) Scatter plot showing correlation (Spearman; r = 0.75; p = 0.026) between pre-CA (baseline) absolute CMRO_2_ and cerebral electrical activity (ECoG IQ) 90 min post-CPR. Blue dots are rats with mild (5 min) ischemia duration, and red dots are rats with prolonged (7 min) ischemia duration. These results suggest that the initial value of CMRO_2_ (pre-injury) may be predictive of cerebral electrical recovery following ischemia and reperfusion.

Fig. 6 shows distributions of CMRO_2_ values measured with our imaging setup as compared to values reported in the literature using various medical imaging approaches, including magnetic resonance methods (MRI/MRS) [23-25] and positron emission tomography (PET) [1-5]. The LSI+SFDI method reported here measures absolute CMRO_2_ values that are within the range measured with these approaches, suggesting the accuracy of our optical imaging approach to determine CMRO_2_.

**Fig. 6.**
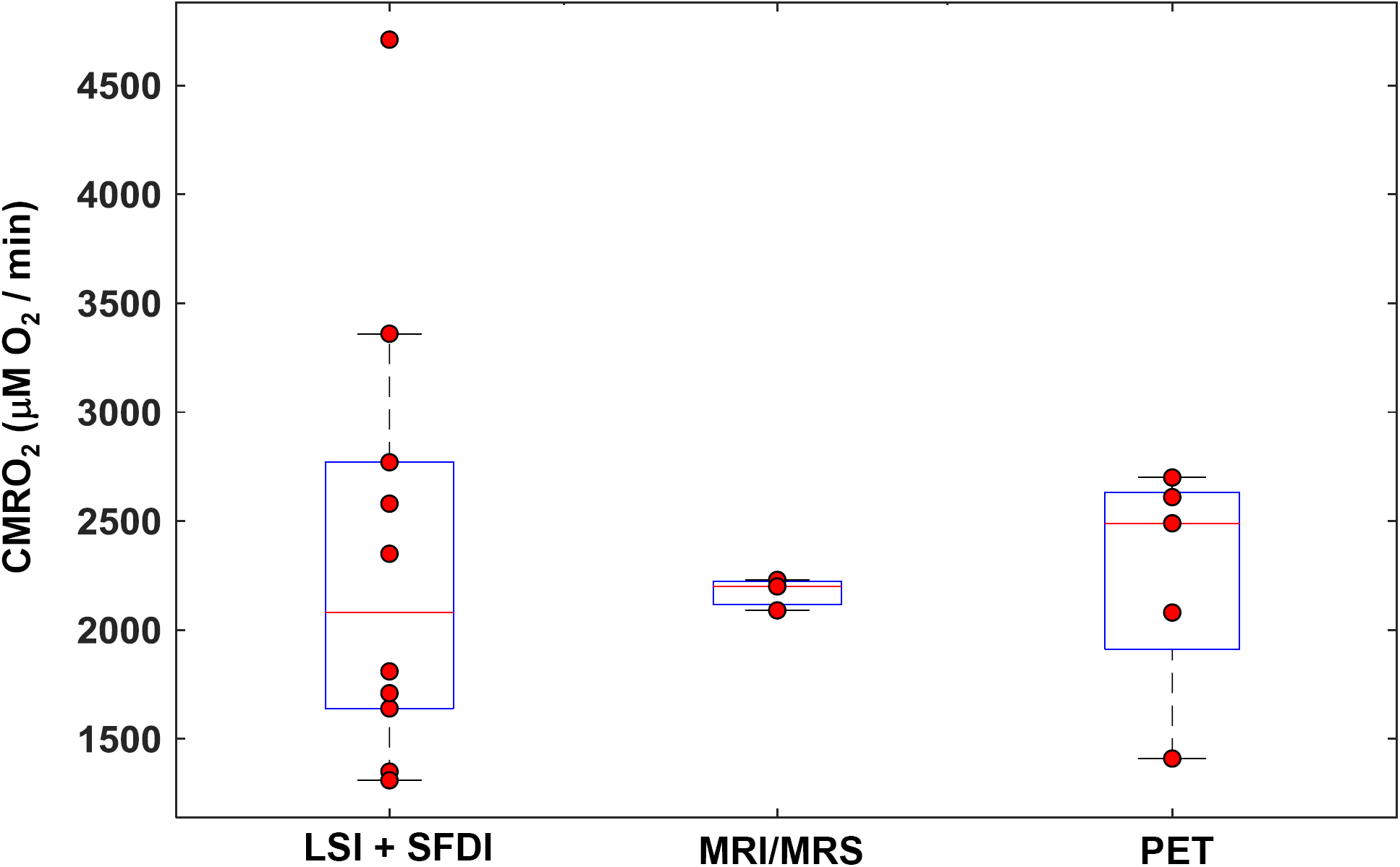
Comparison of absolute CMRO_2_ values (μM O_2_ / min) from our combined SFDI + LSI optical system with absolute CMRO_2_ values reported in other published studies using different imaging modalities (MRI/MRS; [23-25], PET; [1-5]). Each LSI + SFDI data point represents an individual rat (n = 10) imaged in this report. Each data point for MRI/MRS and PET represents the average over all of the rats measured in a separate individual study. Good overall agreement is observed between absolute CMRO_2_ values obtained with our optical imaging technique and those measured with other common imaging modalities.

## Discussion

In this report, we provide, to the best of our knowledge, the first demonstration of dynamic imaging of absolute cerebral metabolic rate of oxygen (CMRO_2_) in the living brain using a combination of laser speckle and spatial frequency domain techniques. We use the tissue absorption and reduced scattering coefficients measured with SFDI to account for the effects of optical properties on our interpretation of the LSI information. We then use the “zero-flow” condition inherent in our CA experimental paradigm to solve for the coefficient α in the CMRO_2_ equation by using a continuity condition at the boundary between normal flow and zero-flow states. Using this technique, we perform quantitative spatial mapping of absolute CMRO_2_ continuously throughout the different stages of the CA + CPR experiment. The CMRO_2_ obtained from our optical system agreed well with established brain imaging techniques (PET, MRI/MRS). Furthermore, we show a significant correlation between absolute CMRO_2_ values at baseline, and longer-term cerebral electrical recovery (ECoG IQ 90 min post-CPR), regardless of CA duration.

This paradigm for measuring CMRO_2_ removes the long-standing requirement that a baseline measurement is acquired, as the CMRO_2_ is now on an absolute quantitative scale (in units of μM O_2_/min) and can thus be directly compared between animals or across multiple separate imaging sessions on different days for a single animal. This enables longitudinal monitoring of cerebral recovery for days or weeks following ischemia and reperfusion without need for a baseline and perturbation (e.g., a gas challenge) on each measurement day. The methods described here are potentially widely applicable to quantitative measurement of metabolic recovery and flow-metabolic coupling and uncoupling in preclinical models of ischemic conditions such as CA and stroke.

### Optical Imaging Segments Venous Regions to Better Quantify Cerebral Oxygen Extraction

The imaging capability of our device allows the segmentation of a region of interest atop a prominent vein, which enables more accurate measurements of the quantity of deoxygenated venous blood and, hence, the quantity of oxygen consumed by the brain. With the use of a larger region of interest, the local CMRO_2_ would be systematically underestimated due to inclusion of the parenchyma in the region; oxygen extraction in the parenchyma is lower than in individual vessels. CMRO_2_ models of diffuse light transport implicitly assume that the concentration of deoxygenated hemoglobin is that within the veins specifically, and not the bulk tissue [26]. However, most diffuse-optics CMRO_2_ measurements are practically unable to satisfy this condition, as they typically use fiber-based spectroscopic techniques that sample the bulk tissue and thus cannot distinguish between venous and mixed arterial-venous parenchymal regions. In this report, the use of diffuse optical imaging allows us to use deoxyhemoglobin concentrations measured in a venous ROI to overcome this limitation and thus obtain more accurate quantitative values of CMRO_2_.

### Correction of CMRO_2_ Data for Partial-Volume Effects

Furthermore, most diffuse-optics measurements of absolute CMRO_2_ are hampered by the partial-volume effect, as the CMRO_2_ equation requires a measurement of the concentration of deoxyhemoglobin *within the vein itself*, not within the tissue as a whole. Therefore, typical diffuse-optics measurements of hemoglobin concentrations in bulk tissue are roughly two orders of magnitude lower than the concentration of hemoglobin in the vein itself. Here, we overcome this limitation by introducing a partial-volume correction to the CMRO_2_ equation. This correction allows us to convert the bulk tissue hemoglobin concentration into an *intravascular* hemoglobin concentration. Without this correction factor, the CMRO_2_ values would also be underestimated by two orders of magnitude. To accurately incorporate this scaling term, it is necessary to know the concentration of total hemoglobin (Hb_bl_) within the blood of each animal. In this report, these values were acquired via arterial blood gas (ABG) measurement before CA. There was a non-negligible amount of variation (mean = 11.95 g/dL; standard deviation = 1.55 g/dL) among the Hb_bl_ values of the 10 subjects in this study. Therefore, failing to account for the variation in Hb_bl_ between the different rats could introduce an additional error of ∼12-25% in the measured CMRO_2_ due specifically to within-group variability in Hb_bl_ values.

### Contributions of Directed Flow versus Diffuse Flow

It is important to note that when we solved for the characteristic flow speed in this report, we assumed that all of the corrected flow speed could be attributed to a “directed-flow” term (Eq. 2). Previous studies have used a Brownian diffusion term as the free parameter when fitting for flow speed [27] or constrained the fit in a model system such that one could choose to fit for either diffuse or directed flow but not both simultaneously [20]. Recently, Postnov et al. [28] used high-speed LSI to map the autocorrelation function pixel-by-pixel in the rodent brain, identifying the dominant type of particle motion at each pixel. In that study, the directed flow term was dominant in large vessels, while the diffuse-motion term was dominant in the parenchyma.

Here, we could not rigorously solve for the autocorrelation function because the sampling frequency of our LSI data acquisition was too low to perform a method similar to that of Postnov et al. [28]. Instead, we used a two-step approach of (1) using SFDI data to account for the effects of optical properties on interpreting the LSI data, and (2) fitting the resulting corrected data to a model of directed flow to extract the characteristic flow speed. This method provided characteristic flow speeds that were similar to previously-reported values [29].

To examine the effect of including both diffuse and directed motion as free parameters in a single model, we used an alternate algorithm (Fig. 7) in which the absorption- and static-scatter-corrected CBF data were fit with a linear combination of both diffuse flow and directed flow, to determine the relative contribution of each type of motion at each pixel. This model extracted a diffusion coefficient that was roughly homogeneous over the entire imaged region (1.4×10^−5^ ± 0.7×10^−5^ mm^2^/s in large vessel ROI at baseline; 1.5×10^−5^ ± 0.8×10^−5^ mm^2^/s in parenchyma ROI at baseline). Also, this model extracted directed flow speeds that were higher in the large vessels (0.78 ± 0.05 mm/s in large vessel ROI at baseline) than in the parenchyma (0.45 ± 0.07 mm/s in parenchyma ROI at baseline). These results support the finding of Postnov et al. [28] that the directed-flow term dominates in the large vessels and the diffuse-flow term contributes more notably in the parenchyma.

**Fig 7.**
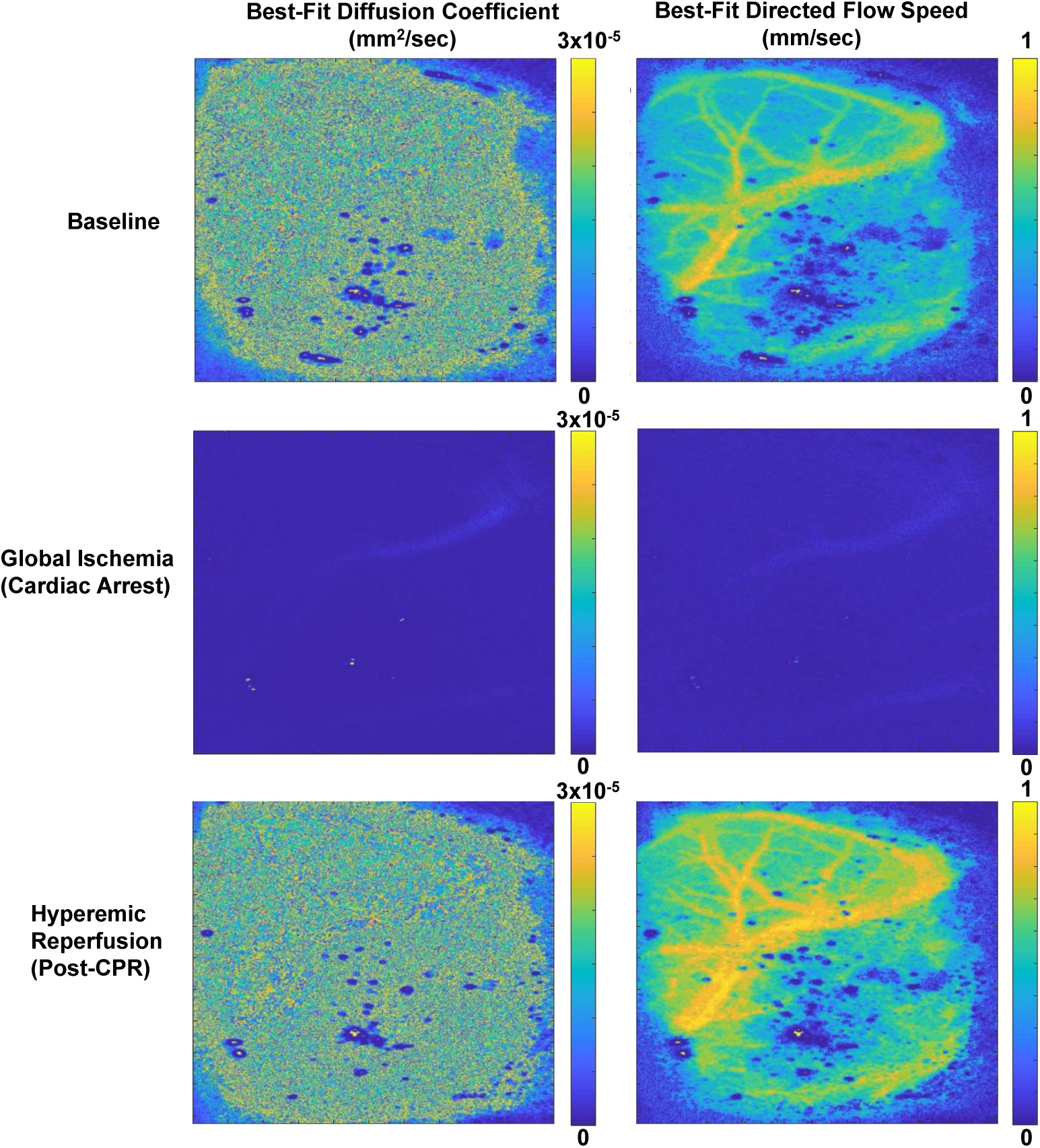
Diffuse and directed flow maps obtained from the brain of a representative rat at baseline, global ischemia (CA), and hyperemic reperfusion post-CPR using a two-term model (one diffusion term, one directed-flow term) to fit to the optical property-corrected LSI data. As expected, during the periods when the brain is perfused, the directed-flow term is high in the large vessels and low in the parenchyma, while the diffusion coefficient is homogeneous in distribution. During CA, the directed-flow term vanishes nearly completely, and the diffusion coefficient is reduced to values more similar to those in static media.

Both the directed-flow term and the diffuse-flow term were considerably reduced during ischemia (CA). The extracted diffusion coefficient and directed flow speed obtained with this two-term model were both similar to previously-reported values [27, 29, 30]. When the directed-flow term alone was used in the model, the extracted characteristic flow speeds were 0.83 ± 0.04 mm/s in the large vessel ROI and 0.52 ± 0.03 mm/s in the parenchyma ROI for the rat shown in Fig. 7. Therefore, using only directed flow in the model is expected to introduce only a small discrepancy (∼10%) in the extracted flow speed, relative to using the two-term approach.

### Limitations of Zero-Flow Condition

Our current approach for measuring absolute CMRO_2_ requires temporary induction of a “zero-flow” condition in the brain. In this report, this condition was met by using a CA model in rats. However, there is a clear need to investigate alternative approaches for interrogating absolute CMRO_2_ without creating harmful perturbations. Future work can incorporate techniques such as temporarily clamping the middle cerebral artery [31] or administering sub-lethal doses of potassium chloride [32] to temporarily induce a zero-flow condition that can be quickly reversed without long-term harm to the animal. It is important to note that once this procedure is performed once on a given animal, subsequent measurements on that same animal should not require additional instances of that procedure. Although the comparison of absolute CMRO_2_ calculated with our approach with PET and MRI is encouraging, further comparison work is required with measurements collected from the same animals under identical anesthesia conditions. Future work is warranted.

## Conclusion

In this report, we have described, to the best of our knowledge, the first mapping of the absolute cerebral metabolic rate of oxygen (CMRO_2_) in the rat brain using diffuse optical imaging. The CMRO_2_ allows for quantitative assessment of the cerebral metabolism without the need for baseline measurements, enabling longitudinal comparison between animals and between multiple days of measurement on an absolute scale. The CMRO_2_ measurements provided by our multimodal system were in good agreement with those previously measured in the brain of anesthetized rats using PET and MRI. This method shows significant potential for assessing and monitoring cerebral metabolism and predicting cerebral response to ischemic injury.

## Acknowledgements

This work is supported by the Arnold and Mabel Beckman Foundation, the United States National Institutes of Health (P41EB015890), the National Science Foundation Graduate Research Fellowship Program (DGE-1321846, to C.C.), the National Center for Research Resources and National Center for Advancing Translational Sciences, National Institutes of Health (through the following grants: R21EB024793 to Y.A., TL1TR001415-01 to R.H.W., KL2 TR001416 to Y.A., a pilot grant to Y.A., all via UL1 TR001414, and a CTSA pilot grant to Y.A. via UL1 TR001414), and the Roneet Carmell Memorial Endowment Fund to Y.A.. The content is solely the responsibility of the authors and does not necessarily represent the official views of the NIH.

